# Motor learning decline with age is related to differences in the explicit memory system

**DOI:** 10.1101/353870

**Authors:** Noham Wolpe, James N. Ingram, Kamen A. Tsvetanov, Richard N. Henson, Rogier A. Kievit, Daniel M. Wolpert, James B. Rowe, for Cambridge Centre for Ageing and Neuroscience

## Abstract

The ability to adapt one’s movements to changes in the environment is fundamental in everyday life, but this ability changes across the lifespan. Although often regarded as an ‘implicit’ process, recent research has also linked motor adaptation with ‘explicit’ learning processes. To understand how these processes contribute to differences in motor adaptation with age, we combined a visuomotor learning paradigm with cognitive tasks that measure implicit and explicit processes, and structural brain imaging. In a large population-based cohort from the Cambridge Centre for Ageing and Neuroscience (n=322, aged 18-89 years) we first confirmed that the degree of adaptation to an angular perturbation of visual feedback declined with age. There were no associations between adaptation and sensory attenuation, which has been previously hypothesised to contribute to implicit motor learning. However, interactions between age and scores on two independent memory tasks showed that explicit memory performance was a progressively stronger determinant of motor learning with age. Similarly, interactions between age and grey matter volume in the medial temporal lobe, amygdala and hippocampus showed that grey matter volume in these regions became a stronger determinant of adaptation in older adults. The convergent behavioural and structural imaging results suggest that age-related differences in the explicit memory system is a contributor to the decline in motor adaptation in older age. These results may reflect the more general compensatory reliance on cognitive strategies to maintain motor performance with age.

**SIGNIFICANCE STATEMENT:** The central nervous system has a remarkable capacity to learn new motor skills and adapt to new environmental dynamics. This capacity is impaired with age, and in many brain disorders. We find that explicit memory performance and its associated medial temporal brain regions deteriorate with age, but the association between this brain system and individual differences in motor learning becomes stronger in older adults. We propose that these results reflect an increased reliance on cognition in order to maintain adaptive motor skill performance. This difference in learning strategy has implications for interventions to improve motor skills in older adults.

## INTRODUCTION

The sensorimotor system has a remarkable capacity to adapt to changes that occur both externally in the environment and internally in neuronal and musculoskeletal dynamics. Such adaptation is critical for learning new skills, and for adjusting previously learned movements in the face of new tasks (Scott, 2004; Franklin and Wolpert, 2011; Wolpert et al., 2011). For example, developmental and ageing processes that occur throughout the lifespan ‒ from changes in muscle and joint physiology to neuronal degeneration in the nervous system ‒ require constant adaptation. However, motor adaptation itself is impaired with age (Fernández-Ruiz et al., 2000; Buch et al., 2003; Seidler, 2007; King et al., 2013), which may put older people at increased risk of adverse events, such as falls (Tinetti et al., 1988; Rubenstein, 2006).

To understand the effects of age on motor learning, optimal control theory proposes that during the execution of a voluntary movement, the central nervous system continuously simulates one’s interaction with the environment (for a review see Franklin and Wolpert, 2011). This may be achieved through an internal forward model, which learns to predict the sensory outcome of an action (Miall and Wolpert, 1996). An error signal between the predicted and actual sensory information leads to the update of the internal model, which facilitates better prediction and improved performance of future actions (Shadmehr et al., 2010). Updating an internal model is believed to be an implicit learning process, central to motor adaptation (Shadmehr et al., 2010; Wolpert et al., 2011).

We previously suggested that this implicit process would be impaired in older adults as a result of reduced reliance on sensory feedback during movement with age (Wolpe et al., 2016). Typically, a reduction in the precision of sensory afferents relative to predictive signals during movement leads to sensory attenuation (Bays et al., 2006). This attenuation is increased with age, with reduced precision of sensory signals and increased reliance on established internal models for motor control (Wolpe et al., 2016). Since the updating of one’s model depends on the relative precision of prediction and sensory signals (Wolpert et al., 2011), the imprecise sensory signals that occur with age would be less able to update the internal model (Lei and Wang, 2017).

Although motor adaptation was once considered to be an archetype of implicit memory, an additional *explicit* learning process has been shown to contribute to motor adaptation (Mazzoni and Krakauer, 2006; Taylor and Ivry, 2011). This explicit process is proposed to be supported by high-level cognitive strategies that counteract changes in the environment (Taylor and Ivry, 2013). On this basis, reduced adaptation could result in part from the age-related decline in the explicit (declarative) memory system (Langan and Seidler, 2011; Trewartha et al., 2014).

Here we test the hypothesis that differences in sensory attenuation and explicit memory contribute to the decline in motor adaptation with age. Participants were recruited from a large population-derived cohort, aged 18-89 years, at the Cambridge Centre for Ageing and Neuroscience (Cam-CAN; Shafto et al., 2014). Participants performed a visuomotor rotation learning task (c.f. Buch et al., 2003), in which they moved a stylus-controlled cursor to a visual target. We then introduced a 30° angular rotation of visual feedback between the cursor and stylus location. Participants therefore needed to adapt their movement to overcome this visuomotor rotation so as to reach the target. We hypothesised that reduced adaptation with age is related to differences in both implicit and explicit processes, including sensory attenuation and declarative memory performance. In addition, we performed whole-brain analyses of grey matter volume to test the corollary hypothesis that differences in adaptation with age are differentially related to grey matter in regions associated with explicit learning, including the hippocampus and amygdala (Hamann et al., 2014; Mary et al., 2017); and regions associated with implicit motor learning, such as the cerebellum, striatum and motor cortex (Seidler et al., 2006; Galea et al., 2011; King et al., 2013).

## MATERIALS AND METHODS

### Experimental design

From a population-based cohort of healthy adults, 322 participants completed a visuomotor learning task (Cam-CAN; Shafto et al., 2014). They were asked to move a cursor so as to hit a target (Fig. 1A). To do so, they grasped a stylus pen with their dominant hand, and the position of the tip of the stylus was recorded using a digitising touch pad (Bamboo CTH-661, Wacom Technology Corporation, Vancouver, WA) and displayed as a red cursor (radius 0.25 cm) on a computer monitor. Participants viewed the display in a semi-reflective mirror, such that the image appeared to be projected onto the horizontal surface of the touch pad. In this way, the red cursor could track the position of the stylus on the pad. The task was to move the cursor from a central ‘home’ position (white disc radius 0.5 cm) to hit one of four possible targets (yellow discs, radius 0.5 cm). Targets were displayed 5 cm from the home position and target direction was chosen from the set {0, 90, 180, 270°}, in a pseudo-random order, such that each cycle of 4 trials contained each target direction. When participants successfully hit a target, it burst and a tone was played to indicate that the trial was successful. If participants failed to initiate movement within 1 sec; or to hit the target within 800 ms after movement initiation, an error tone was played and the message “Too slow” was displayed. Participants completed an initial familiarisation phase of 24 trials (6 cycles of the 4 targets), during which they were permitted to see their hand and the stylus through the mirror. In the main experiment, an occluder was placed behind the mirror to prevent participants from seeing their hand.

**Figure 1.**
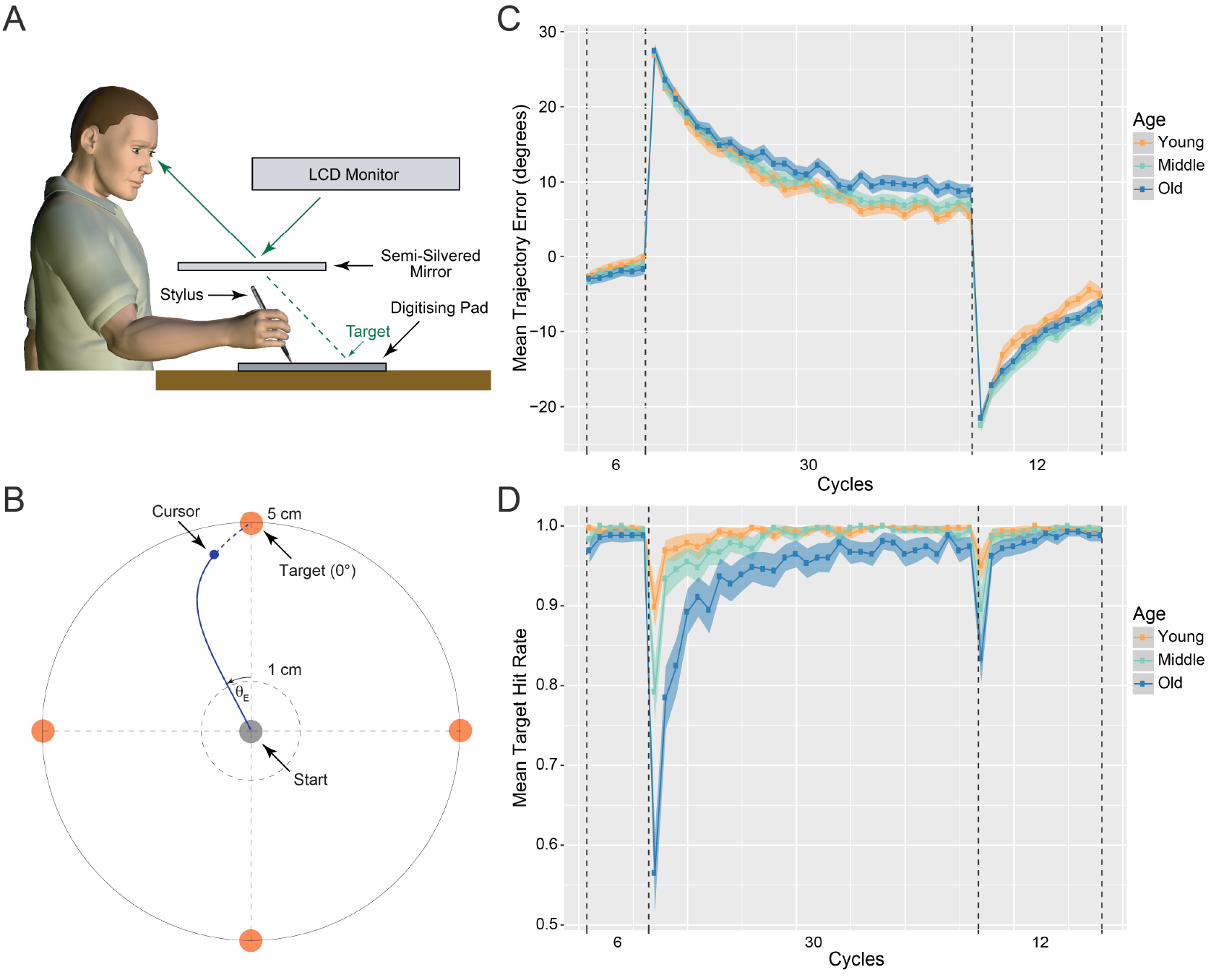
Visuomotor rotation learning task. A. Illustration of the task in which participants moved a stylus-controlled cursor so as to hit a target. The target appeared pseudorandomly in one of four locations on the screen (once in each of the four-trial cycles). Participants could not see their hand, and the visual feedback of the cursor was either veridical (pre-exposure and post-exposure phases) or rotated by 30 degrees (exposure phase) relative to the stylus. B. Participant movement adaptation was assessed by looking at the changes in their initial trajectory error θ_E_, calculated 1 cm after starting the movement C. Mean trajectory error across the experimental cycles (±1 standard error shaded). Dashed vertical lines separate the phases: pre-exposure (left), exposure (middle) and post-exposure (right). For illustration purposes only, data has been split into three age groups of similar size (‘young’ = 18-45 years, N=106; ‘middle’ = 41-65 years, N=106; ‘old’ = 66-89 years, N=107), although all analyses were performed with age as a continuous variable. D. Same ac (C) but for target hit rate.

The main experiment consisted of 192 trials which were divided into three phases. During the pre-exposure phase, participants performed 24 trials (six cycles of four trials) in which the red cursor accurately represented the position of the stylus. During the subsequent exposure phase, participants performed 120 trials (30 cycles) in which the position of the cursor was rotated 30° clockwise relative to the central home position. The introduction of the rotation required participants to adapt their movement trajectories in order to successfully hit the targets. Finally, during the post-exposure phase, participants performed 48 trials (12 cycles) with the perturbation removed, as in the pre-exposure phase. The post-exposure phase required participants to ‘de-adapt’ their movement trajectories in order to hit the target.

Participants also completed two additional tasks that measure processes relevant for implicit and explicit learning: 1) A Force Matching task, measuring sensory attenuation as a proxy of the precision of forward models (n=311 complete datasets) (Wolpe et al., 2016); 2) a Story Recall task, which is a verbal memory subtest of the Wechsler Memory Scale measuring explicit memory (Shafto et al., 2014) (n=319). A smaller subset of the participants (n=116) also completed an Emotional Memory task, which had many more trials and so potentially provides a more sensitive measure of explicit memory (Henson et al., 2016).

### Behavioural statistical analysis

Motor adaptation on each trial was assessed by measuring the initial movement trajectory error, which is considered to reflect the feedforward component of the movement, before feedback becomes available. The trajectory error was calculated as the difference between the target angle and the angle of the initial cursor movement trajectory. The initial trajectory angle was calculated at 1 cm into the movement, relative to the start position (trials were excluded if the cursor moved less than 1 cm from the home position, affecting 0.76% of trials on average across participants). Trajectory errors were averaged across each cycle of 4 trials to give a time series across the 48 cycles (from 192 trials) of the experiment.

For each participant, trajectory errors across cycles in the exposure and post-exposure phases were each fit with an exponential. The fitting algorithm (‘nlinfit’ function in Matlab 2017a; MathWorks Inc. MA, USA) used iteratively reweighted least squares with a bisquare weighting function. The curves were constrained as follows: the exponential for the exposure phase started at 30° on cycle 1 with a variable final value on cycle 30. For the post-exposure phase, the initial value on post-exposure cycle 1 is constrained by the final level of exposure phase adaptation (exposure cycle 30) and asymptote at zero.

The fitting therefore had three free parameters: 1) Final adaptation (in degrees), which is the difference between the angular perturbation of 30° and the fitted trajectory error on the last cycle of the exposure phase (between 0° and 30°); 2) exponential time constant for adaptation (in trials); 3) de-adaptation time constant (in trials). Based on the fit, we also calculated: 1) final de-adaptation, which is the trajectory error on the last cycle of the post-exposure phase. 2) Time to half adaptation, which is the time (in cycles) to reach half the final adaptation; 3) Time to half de-adaptation (in cycles). Time to half adaptation and de-adaption were chosen for the analyses as they were more robust across subjects compared to the exponential time constants. Three participants (aged 28, 48 and 58 years old) were excluded because their fitted final adaptation was 0 degrees, implying failure to understand or perform the task (> 5 SD from cohort mean). De-adaptation was assessed as the absolute ratio between final de-adaptation and final adaptation.

To examine the contribution of processes supporting implicit and explicit learning to age-related differences in motor adaptation, the data were entered into linear regression models. Final adaptation was the dependent variable, and the independent variables were: 1) sensory attenuation, measured as the overall mean force overcompensation when directly matching the target forces (Wolpe et al., 2016); 2) explicit memory in the Story Recall task, measured as the first principal component of the scores given by the experimenter for retelling the story (i) immediately and (ii) 30-minutes after hearing the story (Shafto et al., 2014). An additional exploratory analysis was performed using declarative memory score from the Emotional Memory task. This score was measured as the first principal component of (i) the total number of detail correct background pictures, and (ii) the total number of detail and gist correct pictures ‒ both measures collapsed across emotional valence (Henson et al., 2016). Covariates of no interest included mean trajectory error during the pre-exposure phase (accounting for individual movement bias, e.g. see Buch et al., 2003), education (categories according to Table 1), gender (categorical variable) and handedness (Edinburgh Handedness Score; Oldfield, 1971). All variables were z-scored before entering the regression analysis. Multiple regression was performed as a path model using the Lavaan package (Rosseel, 2012) in R (R Core Team, 2016), using Full Information Maximum Likelihood to account for missing data. All statistical analyses were performed with a two-tailed alpha threshold of 0.05, but given the large sample size, we focus on effect size, here reported as the percentage of variance explained by the specific statistical contrast (R^2^; values less than ~1.2% correspond to two-tailed *p* > 0.05). For the regression analyses, we report the raw as well as fully standardised path estimates. Plots were generated using ggplot2 (Wickham, 2009).

**Table 1.** Summary of participant demographics across age decades.

| Age | N | Gender | Handedness | Education* |  |  |  |
| --- | --- | --- | --- | --- | --- | --- | --- |
|  |  | M/F | R/L | None | GCSE | A Levels | University |
| 18-29 | 33 | 13/20 | 30/3 | 0 | 5 | 6 | 22 |
| 30-39 | 46 | 24/22 | 40/6 | 0 | 2 | 7 | 37 |
| 40-49 | 60 | 28/32 | 51/9 | 1 | 7 | 4 | 48 |
| 50-59 | 46 | 25/21 | 41/5 | 3 | 5 | 15 | 23 |
| 60-69 | 55 | 31/24 | 50/5 | 3 | 10 | 14 | 28 |
| 70-79 | 53 | 22/31 | 49/4 | 6 | 8 | 9 | 30 |
| 80-89 | 29 | 16/13 | 28/1 | 5 | 4 | 12 | 8 |
| Total | 322 | 159/163 | 289/33 | 18 | 41 | 67 | 196 |

\*Categorised according to the British education system: 'none' = no education over the age of 16 years; 'GCSE' = General Certificate of Secondary Education; 'A Levels' = General Certificate of Education Advanced Level; 'University' = Undergraduate or graduate degree.

### Structural neuroimaging protocol and analysis

A 3T Siemens TIM Trio with a 32-channel head coil was used to scan 310 participants (12 participants declined MRI). Both a T1-weighted MPRAGE image (TR 2250 ms, TE 2.99 ms, TI 900 ms, FA 9°, FOV 256 mm × 240 mm × 192 mm, isotropic 1 mm voxels) and a T2-weighted SPACE image (TR 2800 ms, TE 408 ms, FOV 256mm × 256mm × 192mm; isotropic 1mm voxels) were acquired. The MR data of eight participants were not included in the analysis due to technical problems during scanning or preprocessing problems. Together with the exclusion of three participants due to outlying behavioural data (see above), 299 participants were included in the structural imaging analyses.

The structural images were preprocessed for a Voxel-Based Morphometry analysis, as previously described (Taylor et al., 2017) using SPM12 (www.fil.ion.ucl.ac.uk/spm) as called by the automatic analysis batching system (Cusack et al., 2015). Multimodal segmentation (using both T1- and T2-weighted images) was used to reduce age-biased tissue priors. Diffeomorphic Anatomical Registration Through Exponentiated Lie Algebra (DARTEL) approach was applied to improve inter-subject alignment (Ashburner, 2007) as follows: segmented images were warped to a project-specific template, and then affine-transformed to the Montreal Neurological Institute (MNI) space, followed by modulation by the Jacobean of the combined transformations (to preserve volume) and smoothing with an 8-mm full-width at half maximum Gaussian kernel. A threshold of 0.15 was used on these images for the inclusion of grey matter voxels, as in previous analysis (Wolpe et al., 2016). Multiple regression analysis was performed to create a statistical parametric map of differences in grey matter volume in relation to adaptation. Adaptation and the (mean-corrected and orthogonalised) interaction term between adaptation and age were included as the main covariates of interest. Age, handedness (Edinburgh handedness score), gender (categorical variable), education (categorical variables according to Table 1), mean pre-exposure trajectory error and total intracranial volume were also included in the regression model. All variables were z-scored before entering the regression analyses. Unless stated otherwise, results of structural imaging analyses are reported at cluster-based *p* < 0.05, Family-Wise-Error (FWE) corrected, with a cluster-forming threshold of *p* < 0.001. The raw data and analysis code are available upon signing a data sharing request form (http://www.mrc-cbu.cam.ac.uk/datasets/camcan/).

## RESULTS

### Differences in motor learning with age

For each participant, we examined the initial movement trajectory error (Fig. 1B) for each cycle across the three experimental phases. Although age was modelled as a continuous variable in all the following analyses, for ease of visualisation, Figure 1C illustrates participants’ trajectory errors for the cohort divided by age into three groups of similar size. During the pre-exposure phase, there was a small but consistent counter clockwise (negative angle) bias in trajectory errors across participants (absolute mean bias less than 2°; *t*_(318)_ = −11.793, *p* = 7.116e-27, *R*^2^ = 0.552). In view of a trend for an effect of age on bias (*t*_(317)_ = −1.933, *p* = 0.054, *R*^2^ = 0.012), we adjusted for individual differences in pre-exposure error in line with previous studies (Buch et al., 2003). In the exposure and post-exposure phases, participants gradually adapted their initial movement to the onset and offset of the 30° angular rotation (Fig. 1C) and improved their performance in terms of target hit rate (Fig. 1D). For the exposure and postexposure phases, we fit the trajectory errors with separate exponential curves (Fig. 2A). The key parameter to assess learning was ‘final adaptation’, i.e. the difference between the 30° angular perturbation and fit trajectory error on the last cycle of the exposure phase. Additional parameters of interest were ‘time to half adaptation’, i.e. the time (in cycles) to reach half the final adaptation; and ‘final de-adaptation’ and ‘time to half deadaptation’ for the post-exposure phase. Across participants, the model fit the data well, with a mean *R*^2^ of 0.742 (*SD* = 0.177), which did not vary with age (*r*(_317_) = −0.100, *p* = 0.076, *R*^2^ = 0.010).

**Figure 2.**
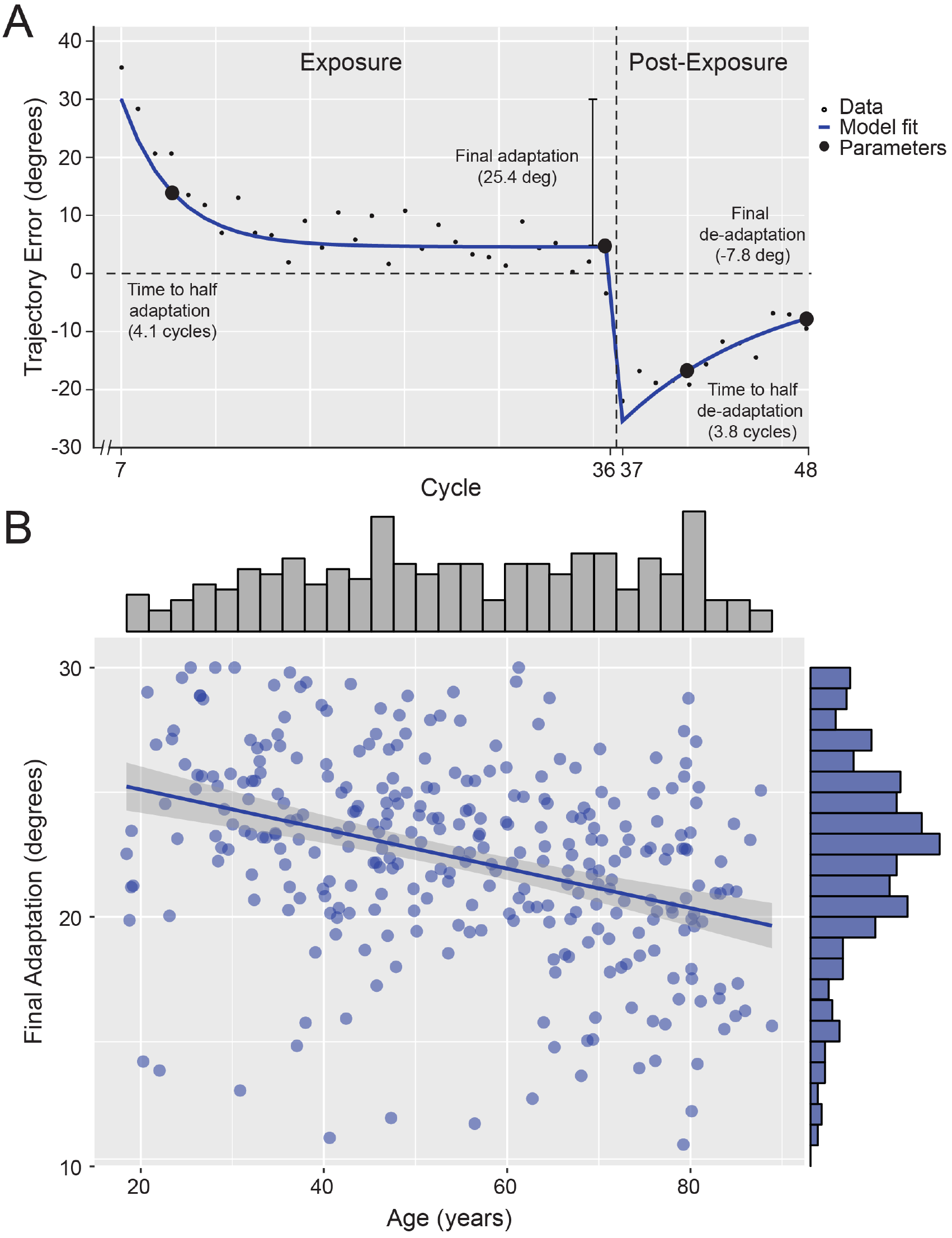
Final adaptation across age. A. Example of the model fit in a representative participant. The model consisted of two sequential exponential curves, fit with a robust bisquare weight function. The main parameter of interest was ‘final adaptation’. B. Correlation between final adaptation and age (with marginal histograms). Solid line indicates the linear regression fit with 95% confidence interval (grey shade).

The magnitude of final adaptation is plotted against age in Figure 2B. We fit the association between final adaptation and age with a linear model (the BIC difference relative to a second-order polynomial model was 2.67 in favour of the linear model). There was a significant negative correlation between age and final adaptation (*r*(_317_) = −0.349, *p* = 1.353e-10, *R*^2^ = 0.122), suggesting that older adults adapted their initial movement trajectory less than young adults. Examining the time-course of individual adaptation, there was a small correlation between ‘time to half adaptation’ and age (*r*(_317_) = −0.1371, *p* = 0.0143, *R*^2^ = 0.019), which became stronger when covarying for final adaptation (partial correlation; *r*(_316_) = −0.201, *p* = 3.101e-04, *R*^2^ = 0.04). Similar results were obtained when examining at the time constant from the exponential fit, which together suggest that although older adults learned less than young adults overall, they did so faster.

In the post-exposure phase, participants ‘de-adapted’ to some degree, but remained biased in the opposite direction to the experimental perturbation (see Figure 1C). Older adults de-adapted less than young adults, with a significant negative correlation between age and final de-adaptation (partial correlation with final adaptation covaried; *r*(_316_) = −0.23, *p* = 3.50e-05, *R*^2^ = 0.053). The time-course for de-adaptation, however, did not vary with age (*r*(_317_) = −0.083, *p* = 0.138, *R*^2^ = 0.007).

### Contribution of implicit and explicit processes to differences in motor learning

To study the potential processes underlying reduced motor adaptation with age, we examined the link between final adaptation and individual differences in measures that could support implicit and explicit learning. We used a measure of sensory attenuation, reflecting the precision of internal models, which may support implicit motor learning (Wolpe et al., 2016), and an explicit measure of declarative memory from a Story Recall task (Shafto et al., 2014). We entered these measures into a linear regression model with final adaptation as the dependent variable, sensory attenuation and declarative memory as the independent variables, as well as their interaction with age, and covariates of no interest (see Methods).

Table 2 summarises the results of the regression analysis. Sensory attenuation was not a significant predictor of adaptation (*beta* = -0.057, *Z* = −1.059, *p* = 0.289, *beta*_standardised_ = -0.057) and there was no age × attenuation interaction (*beta* = 0.04, *Z* = 0.815, *p* = 0.415, *beta*_standardised_ = 0.043; Fig. 3A). Declarative memory also showed no main effect on adaptation (*beta* = -0.04, *Z* = -0.908, *p* = 0.364, *beta*_standardised_ = -0.055), but there was a positive age × declarative memory interaction (*beta* = 0.087, *Z* = 2.123,*p* = 0.034, *beta*_standardised_ = 0.112; Fig. 3B). An analogous interaction was also found with the alternative and exploratory measure of declarative memory, from the Emotional Memory task performed by a subset of participants: again, no main effect of declarative memory was observed (*beta* = 0.093, *Z* = 1.269, *p* = 0.204, *beta*_standardised_ = 0.131), but a positive age × declarative memory interaction emerged (*beta* = 0.171, *Z* = 2.607, *p* = 0.009, *beta*_standardised_ = 0.22). These results suggest that sensory adaptation was more positively correlated with explicit memory in older adults.

**Figure 3.**
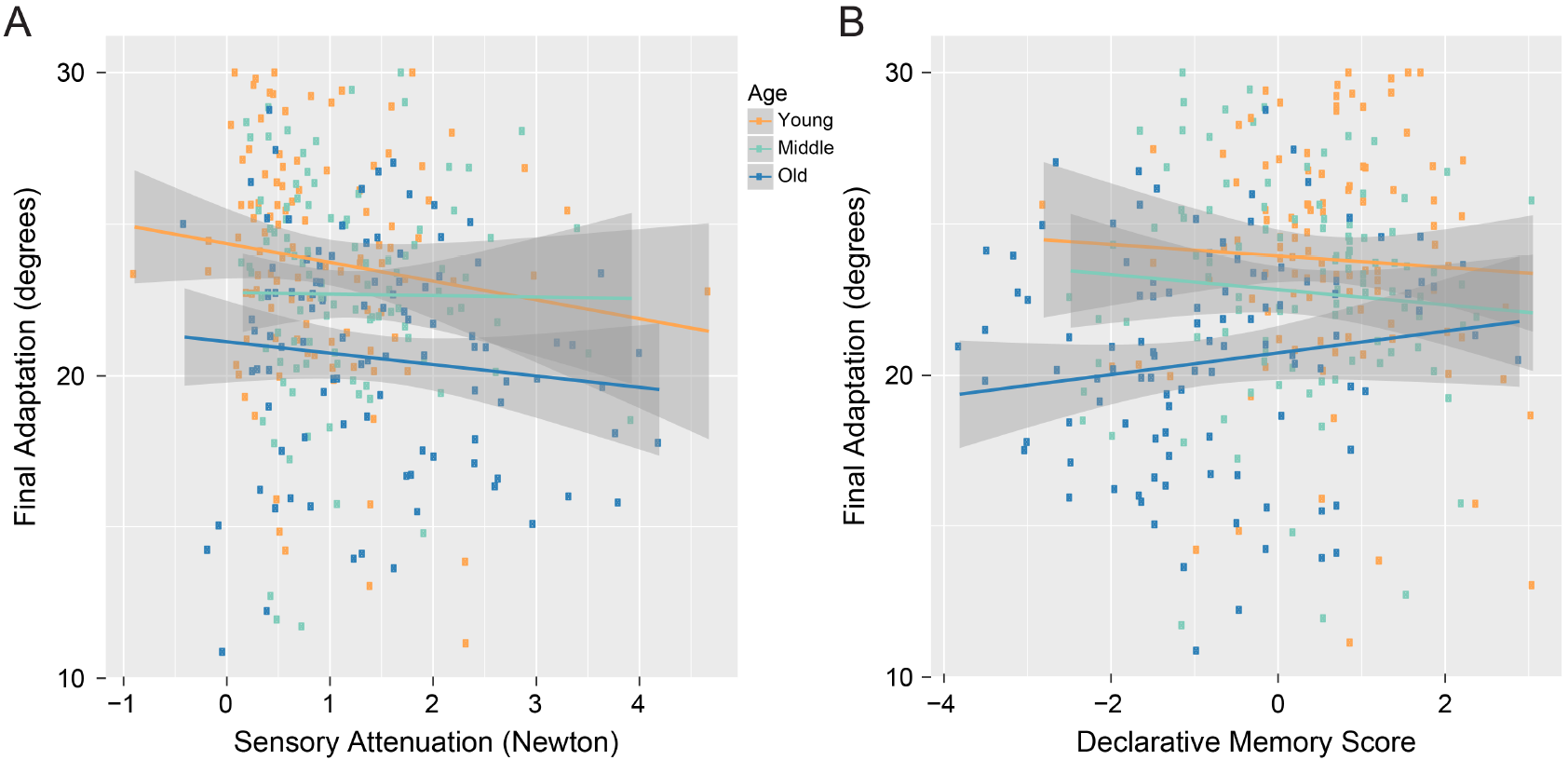
Explicit memory performance and motor adaptation by age. **A.** Illustration of the positive interaction between age and declarative memory scores from the Story Recall task in relation to final adaptation. Age groups as in Figure 1. Solid line indicates the linear regression fit with 95% confidence interval (grey shade). **B.** As in (A), but for declarative memory score from the Emotional Memory task.

**Table 2.**
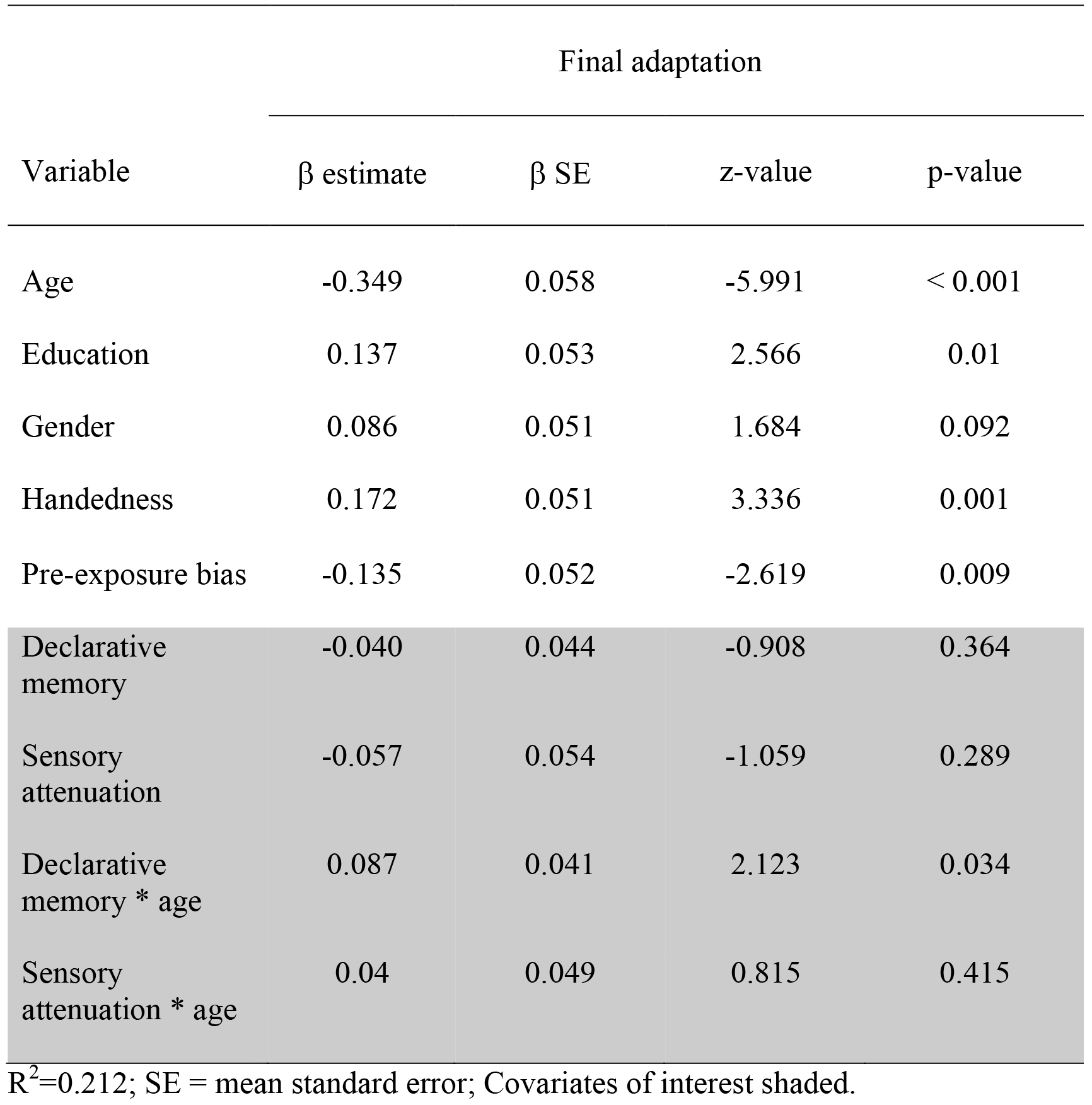
Summary of multiple regression analysis for predicting final adaptation.

### Grey matter differences and reduced adaptation with age

We next performed a spatially-unbiased, whole-brain Voxel-Based Morphometry analysis of grey matter volume. To identify brain areas where grey matter volume was correlated with differences in adaptation across age, we examined the correlation with the interaction of final adaptation × age. There was a significant positive correlation between grey matter volume and adaptation × age in three clusters (Fig. 4A): one encompassing the right middle and inferior temporal lobe (*k* = 1244, *p* = 0.020, *FWE*-corrected) and two bilateral clusters that include the right (*k* =1254, *p* = 0.019, *FWE*- corrected) and left (*k* = 1238, *p* = 0.020, *FWE*-corrected) hippocampus and amygdala. This interaction indicates that grey matter volume in these regions was more positively correlated with final adaptation in older, versus younger participants (Fig. 4B). Given these regions’ involvement in explicit memory, these results are consistent with the behavioural findings implicating a role for explicit memory in age-related differences in motor learning. No significant negative correlation was found with adaptation × age.

**Figure 4.**
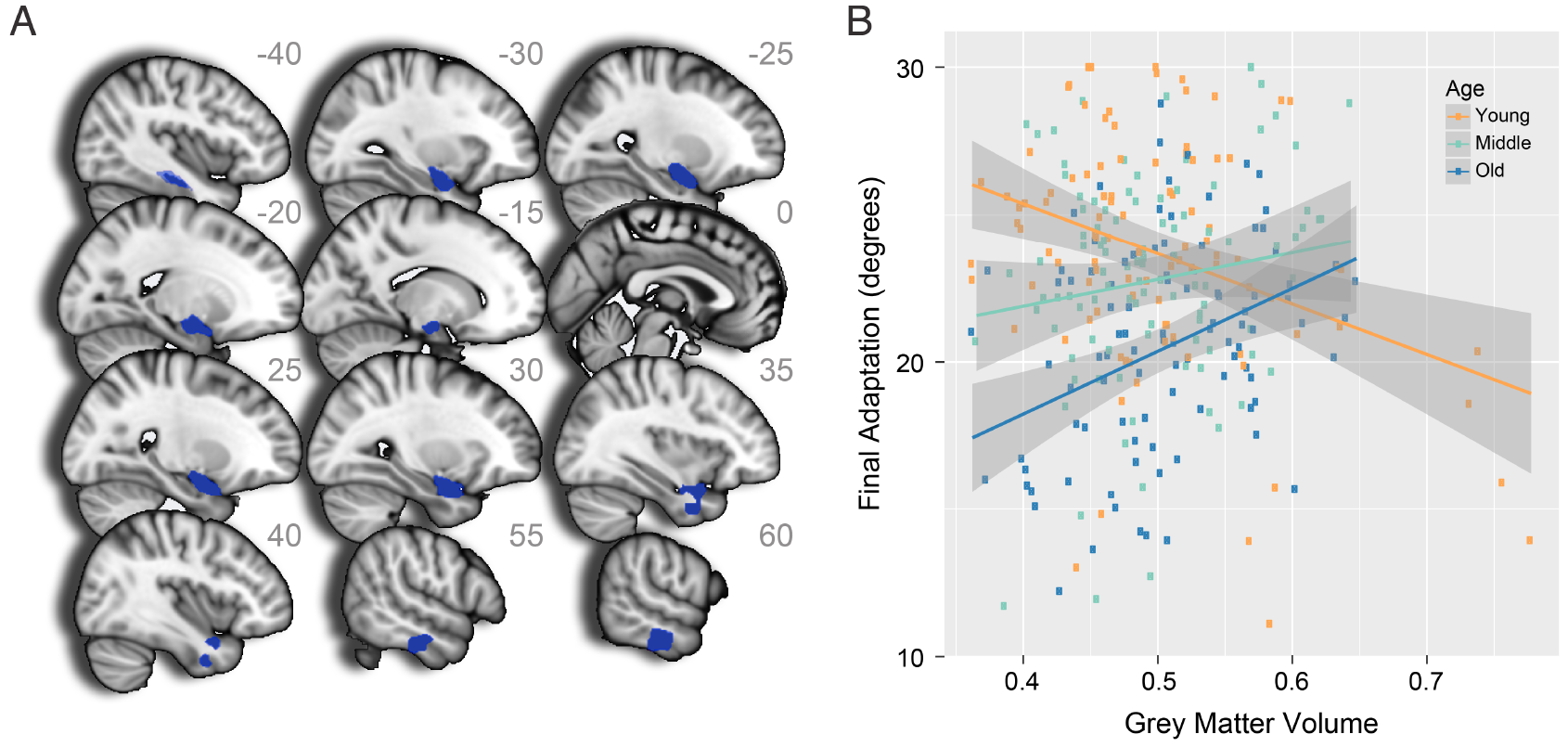
Structural imaging results. A. Sagittal sections (numbers indicating x coordinate), showing three significant clusters (yellow) where there was a significant (p < 0.05, *FWE*-corrected) positive interaction between final adaptation and age in relation to grey matter volume. These clusters included bilateral hippocampus and amygdala as well as right medial and inferior temporal lobe. B. Illustration of the positive interaction from (A). Mean grey matter volume extracted from peak voxel for illustration of interaction. Groups split by age as in Figure 1.

For completeness, we looked at the correlation between grey matter volume and adaptation, independently of age. Here, there was a trend for a positive correlation between adaptation and grey matter volume in the ventral striatum (*k* = 879, *p* = 0.061, *FWE*-corrected; peak voxel in the nucleus accumbens at [-14, −3, −10]). No significant negative correlation was found, however, in a lenient threshold of *p* < 0.001 uncorrected, a small cluster was found in the cerebellum (*k* = 192; peak voxel at [8, −50, −42]).

## DISCUSSION

The results of the current study suggest that the reduction of motor adaptation as we grow older is related to individual differences in explicit memory, but not in sensory attenuation. Across participants, reduced grey matter in brain structures of the medial temporal lobe, including the hippocampus and amygdala, was associated with reduced motor adaptation. These results contrast with the classical view of motor learning as a pure implicit learning process.

### No association between sensory attenuation and motor adaptation

In the classical interpretation of motor learning, an internal forward model predicts the sensory consequences of one’s movement (Shadmehr et al., 2010; Wolpert and Flanagan, 2010). A discrepancy between sensorimotor prediction and feedback (sensory prediction error) enables the internal model to be updated. We hypothesised that this implicit process would contribute to reduced degree of adaptation seen with age in in our study and in previous studies (McNay and Willingham, 1998; Buch et al., 2003; Bock, 2005). Specifically, as ageing leads to reduced reliance on ‘noisy’ sensory information, reflected in increased sensory attenuation (Wolpe et al., 2016), internal models would become progressively less sensitive with age to differences between sensory prediction and feedback.

We found that differences in attenuation did not explain reduced adaptation with age. These findings, coupled with the structural imaging results, suggest that the processes underlying individual differences in adaptation with age differ from those underlying altered sensorimotor integration (Wolpert et al., 2011; Wolpe et al., 2016). Further, the absence of an association between sensory attenuation and motor learning across participants is surprising, considering the theoretical link between these measures (Wolpe et al., 2016). This null result may be because the measure of sensory attenuation reflects the precision-dependent down-weighting of haptic and proprioceptive feedback, whereas our motor adaptation task relied heavily on visual feedback. Attenuation might be related to adaptation in other tasks, with for example a physical force field perturbation, rather than virtual perturbation.

### Contribution of explicit memory to reduced motor learning with age

In recent years, evidence has emerged for the contribution of explicit learning strategies to motor adaptation (Taylor and Ivry, 2012). For example, individual differences in cognitive abilities have been linked with motor adaptation, such as working memory capacity in young (Anguera et al., 2010) and older adults (McNay and Willingham, 1998; Langan and Seidler, 2011; Uresti-Cabrera et al., 2015). In our study, there was an increased association between declarative memory performance and adaptation with age. This suggests that the decline in explicit memory with age (Henson et al., 2016) contributes to reduced motor adaptation. Although the behavioural effect size was not large, a similar effect was observed with another explicit memory task in a subset of our cohort, and builds on three key observations. First, when an experimental perturbation is small and gradual, emphasising implicit processes, older adults adapt their movement as well as young adults (Buch et al., 2003). Second, when young and old participants are matched by explicit knowledge of the perturbation, age-related differences largely dissipate (Heuer and Hegele, 2008). Third, explicit memory performance has been linked to reduced motor learning with age, but specifically in the ‘fast’ learning process (Trewartha et al., 2014).

Rather than a single learning process, a two-state model has been suggested to better explain motor adaptation (Smith et al., 2006), in which there are two learning processes occurring in parallel with a fast and a slow learning rate. The fast learning process has been associated with explicit learning strategies (McDougle et al., 2015), including explicit memory in general (Keisler and Shadmehr, 2010) and in old age in particular (Trewartha et al., 2014). Further, increased awareness to visuomotor perturbations has been linked to an increased early adaptation (Werner et al., 2015). Both our findings that learning was faster in older adults, and the increased association between adaptation and declarative memory in old age, together suggest that older adults rely more on an explicit learning strategy with a fast learning rate.

### Different strategies for motor learning with age

In younger adults, it has been suggested that individuals with better explicit memory rely more on explicit learning during motor adaptation, in order to optimise adaptation capacity (Christou et al., 2016). However, considering the substantial decline in explicit memory with age (Henson et al., 2016), why would older adults rely on a strategy that would lead to their reduced learning? To propose an answer to this question, we consider the mechanisms underlying implicit and explicit learning for motor adaptation.

In contrast to implicit motor learning which is driven by sensory prediction error (see above), the explicit component of motor learning is proposed to be mainly driven by the task performance error ‒ that is, the difference between the target and sensory feedback (Taylor and Ivry, 2013). A careful consideration of movement adaptation and target hit rate (see Figure 1C and Figure 1D) shows that unlike older adults, younger participants continued to adapt their movement trajectories even after performance had reached ceiling level (in terms of successfully reaching the target). In other words, young, but not older adults, continue to adapt in the absence of (explicit) task performance error, possibly owing to the persistence of (implicit) sensory prediction error. This behavioural discrepancy may be due to differential use of ‘cost functions’ with age (Marblestone et al., 2016): whereas younger participants optimise movement in terms of metabolic expenditure, jerk or torque change (Todorov and Jordan, 2002; Todorov, 2004), older adults may be more sensitive to performance error signals (Levy-Tzedek, 2017) in order to maximise immediate task success.

The tighter coupling between task success and motor adaptation in older adults in our study may also speak to increased reliance on a ‘model-free’ strategy for learning. In addition to the ‘model-based’ approach discussed thus far, computational models have described an additional ‘model-free’ approach for learning in general (Daw et al., 2005), and for motor learning in particular (Huang et al., 2011; Izawa and Shadmehr, 2011; Cashaback et al., 2017). In model-free learning, actions are selected so as to maximise the predicted value of reward that is learned through trial and error (Daw et al., 2005). It is computationally efficient, and dependent on dopaminergic signalling of reward prediction error that is distinct from sensory prediction error (Palidis et al., 2018). Model-free learning can indeed account for different behavioural phenomena in motor adaptation, including savings (faster relearning) that is intact in older individuals (Seidler, 2007; Huang et al., 2011). Moreover, a model-free strategy for motor learning is likely to become more prominent when sensory precision is reduced (Izawa and Shadmehr, 2011), as occurs in age (Wolpe et al., 2016; Lei and Wang, 2017). Taken together, model-free learning may remain preserved relative to model-based learning for motor adaptation in old age (c.f. Sharp et al., 2016, but see Chowdhury et al., 2013), however, this remains to be directly tested in future studies.

### Increased association between explicit memory system and motor learning with age

Complementing our behavioural data, bilateral hippocampal and amygdala grey matter volumes were positively associated with adaptation, more so with increasing age. As the medial temporal lobe and hippocampus play a central role in declarative memory, these imaging results underscore the behavioural associations, between explicit memory and motor adaptation through the lifespan. The association is consistent with the notion of increased reliance on cognitive resources in old age for maintaining motor performance (Seidler et al., 2010), e.g. as seen during normal walking (Mirelman et al., 2017). Whether these interactions indeed reflect a (compensatory) behavioural (Seidler and Carson, 2017) and functional (Tsvetanov et al., 2016) reliance on cognition, or simply the larger variability in explicit memory with age remains to be validated.

The anterior part of the hippocampus identified in our study supports the learning of new environmental layouts (Maguire et al., 2000), encoding the Euclidean distance to one’s goal (Howard et al., 2014). This goal distance signal is speculatively analogous to the performance error signal for mediating explicit motor learning (Taylor and Ivry, 2011). Similar performance error signals have been found in the adjacent amygdala (Gemba et al., 1986), which enhances learning of highly arousing or rewarding action-outcome associations (Cador et al., 1989; Fastenrath et al., 2014).

Taken together, the behavioural and imaging results suggest that across the lifespan, adults gradually come to rely more on explicit learning strategies, driven by performance error, in order to maintain success even on a motor adaptation task. Although our study focussed on healthy adults, a gradual increase in the importance of explicit memory for motor learning across the lifespan may also inform the development of more efficient neurorehabilitation programmes at different ages.

## ACKNOWLEDGEMENTS

We are grateful to the Cam-CAN respondents and their primary care teams in Cambridge for their participation in this study. Cam-CAN research was supported by the Biotechnology and Biological Sciences Research Council (BB/H008217/1). JBR and NW were supported by the James S. McDonnell Foundation 21st Century Science Initiative, Scholar Award in Understanding Human Cognition. JBR was supported by Wellcome Trust [103838] and the Medical Research Council [SUAG/004 RG91365]. DW was supported by the Wellcome Trust [097803], Human Frontier Science Program and the Royal Society Noreen Murray Professorship in Neurobiology. RNH was supported by the Medical Research Council [SUAG/010 RG91365]. RAK was supported by the Wellcome trust and the Medical Research Council [SUAG/014 RG91365]. RNH and RAK were also supported by European Union’s Horizon 2020 research and innovation programme under grant agreement No 732592.

